# Quantitative analysis of the ThrbCRM1-centered gene regulatory network

**DOI:** 10.1101/538751

**Authors:** Benjamin Souferi, Mark M. Emerson

## Abstract

Enhancer activity is determined by both the activity and occupancy of transcription factors as well as the specific sequences they bind. Experimental investigation of this dynamic requires the ability to manipulate components of the system, ideally in as close to an in vivo context as possible. Here we use electroporation of plasmid reporters to define critical parameters of a specific cis-regulatory element, ThrbCRM1, during retinal development. ThrbCRM1 is associated with cone photoreceptor genesis and activated in a subset of developing retinal cells that co-express the Otx2 and Onecut1 (OC1) transcription factors. Variation of reporter plasmid concentration was used to generate dose response curves and revealed an effect of binding site availability on the number and strength of cells with reporter activity. Critical sequence elements of the ThrbCRM1 element were defined using both mutagenesis and misexpression of the Otx2 and OC1 transcription factors in the developing retina. Additionally, these experiments suggest that the ThrbCRM1 element is co-regulated by Otx2 and OC1 even under conditions of sub-optimal binding of OC1.

**Summary Statement:** Systematic variation of the levels of a transcriptional reporter plasmid, its trans-acting factors, and transcription factor binding sites reveals properties of a retinal enhancer during development.

## Introduction

The rules and logic of cis-regulatory activity that underlie dynamic gene regulation during development are an area of great interest (Rickels and Shilatifard, 2018). At present, quantitative measurements are largely determined through highly reductionist approaches such as EMSAs or protein microarrays, while in vivo activity is qualitative, limiting the ability to correlate specific sequence elements with reporter output. Elucidation of this process during development is further complicated by temporal dynamics as cells have rapid shifts in active gene regulatory networks (GRNs). However, identification of these networks provides insights into how transcription factor expression and activation are coordinated to direct cell fate choices during development.

Electroporation is one method to identify and characterize cis-regulatory elements as components of GRNs. Several studies have used this method to identify cis-regulatory elements, providing critical insights into retinal development and other developmental contexts (Bery et al., 2014; Emerson et al., 2013; Emerson and Cepko, 2011; Hsiau et al., 2007; Maguire et al., 2018; Mills et al., 2017; Uchikawa et al., 2004; Wang et al., 2014). In addition, the technique of electroporation is widely used to misexpress transcription factors, or other signaling factors (Chang et al., 2013; Cherry et al., 2011; de Melo et al., 2011; Emerson et al., 2013; La Torre et al., 2013; Matsuda and Cepko, 2007; Mattar et al., 2015; Onishi et al., 2010; Wang et al., 2014). However, the effect of specific parameters such as concentration of plasmid DNA are largely unaddressed. To date, it has not been established how reporter or misexpression DNA plasmid concentration affect the output and interpretation of electroporation experiments.

The vertebrate retina is an excellent model organ to investigate the development of nervous system complexity. Recent analysis has suggested that the retina may be composed of as many as 100 cell types, each of which are generated from multipotent retinal progenitor cells during development (Holt et al., 1988; Turner et al., 1990; Wetts and Fraser, 1988; Zeng and Sanes, 2017). Recently, the existence of restricted progenitor cell states that preferentially give rise to certain cell types has been characterized (Bery et al., 2014; Emerson et al., 2013; Emerson and Cepko, 2011; Hsiau et al., 2007; Maguire et al., 2018; Mills et al., 2017; Uchikawa et al., 2004; Wang et al., 2014). One of which gives rise to cone photoreceptors and horizontal cells and can be identified by reporters driven by the Thyroid hormone receptor beta gene cis-regulatory module 1 (ThrbCRM1) element (Emerson et al., 2013). Previous work has suggested that the ThrbCRM1 element is active in retinal progenitor cells that co-express the transcription factors Otx2 and Onecut1 (OC1). Misexpression and loss-of-function analysis supports a model in which Otx2 and OC1 are both required for induction of ThrbCRM1 reporters, likely through direct binding of each transcription factor to the element, though the direct mechanism remains unknown (Emerson et al., 2013).

Here we describe a quantitative analysis of the ThrbCRM1 element in developing retinas. Using electroporation as a method to introduce fluorescent reporter plasmids and flow cytometry to quantitate reporter activity, the activity of ThrbCRM1 elements were measured in terms of total reporter-positive cells as well as fluorescence level of cells within the population. This analysis revealed distinct differences in concentration-dependent reporter activity that depended on the copy number of cis-regulatory elements. Misexpression of the Otx2 and OC1 activating transcription factors also led to concentration-dependent changes in reporter activity that suggested saturation of reporter activation also occurred. Flow cytometry was used to determine the likely functional Otx2 binding site. Lastly, using these mutated ThrbCRM1 plasmids in combination with Otx2 and OC1 misexpression plasmids suggested that the co-requirement for these two transcription factors is likely not at the step of complex formation on DNA.

## Materials and Methods

### Animals

All methods used in animal studies were approved by City College of New York, CUNY animal care protocols. Fertilized chicken eggs and CD-1 mice were obtained from Charles River. Eggs were stored in a 16° room up to 10 days before incubation and incubated in a 38°C humidified incubator. Retinas were isolated from chicken embryos and mouse P0 pups without regard to sex.

### DNA Plasmids

The Stagia3 EGFP reporter plasmid uses a minimal TATA box from Herpes Simplex Virus and is described in (Billings et al., 2010). The co-electroporation plasmids CAG::mCherry (constructed by Takahiko Matsuda and reported in (Wang et al., 2014)), CAG::EGFP (Matsuda and Cepko, 2004), CAG::Nucβ-gal (Emerson and Cepko, 2011), UbiquitinC::TdTomato (UbiqC::TdT) (Rompani and Cepko, 2008) have been described previously. CAG::Otx2 and CAG::OC1 misexpression plasmids use mouse versions of the relevant transcription factors(Emerson et al., 2013; Kim et al., 2008). The Thrb reporters ThrbCRM1(2X)::EGFP and ThrbCRM1(4X)::EGFP used in Figures 1, 2, 3 and 4 were previously reported (Emerson et al., 2013). The ThrbCRM(2X)::EGFP reporter used in Figure 5 differs from the reporter used in previous Figures with regards to the restriction enzyme sites used to insert the 40 base pair ThrbCRM1 element into Stagia3. Briefly, one pair of complementary ThrbCRM1 oligos were designed such that annealing produced a double-stranded DNA with a Sal1 overhang on one end and a Hind3 overhang on the other. A separate pair of complementary ThrbCRM1 oligos were designed such that annealing produced a double-stranded DNA molecule with a Hind3 overhang on one end and an EcoR1 overhang on the other end. Oligo pairs were annealed and phosphorylated by T4 Polynucleotide kinase enzyme (NEB, M0201S), chloroform extracted, and precipitated overnight. A triple ligation (Takara, 6022) with Stagia3 digested with Sal1 and EcoR1 restriction enzymes produced clones with two copies of the ThrbCRM1 element oriented the same way in Stagia3 and joined by a Hind3 restriction site. Oligos encoding the mutant forms of ThrbCRM1 described in Figure 5 were cloned in a similar manner to the wildtype oligos. All constructs were verified by Sanger sequencing. All DNA plasmids used in electroporation experiments were purified using Midiprep DNA isolation kits (Qiagen, 12143) and resuspended in Tris-EDTA (TE) buffer. DNA concentration and purity was verified using a Nanodrop 1000 (Thermoscientific).

**Figure 1.**
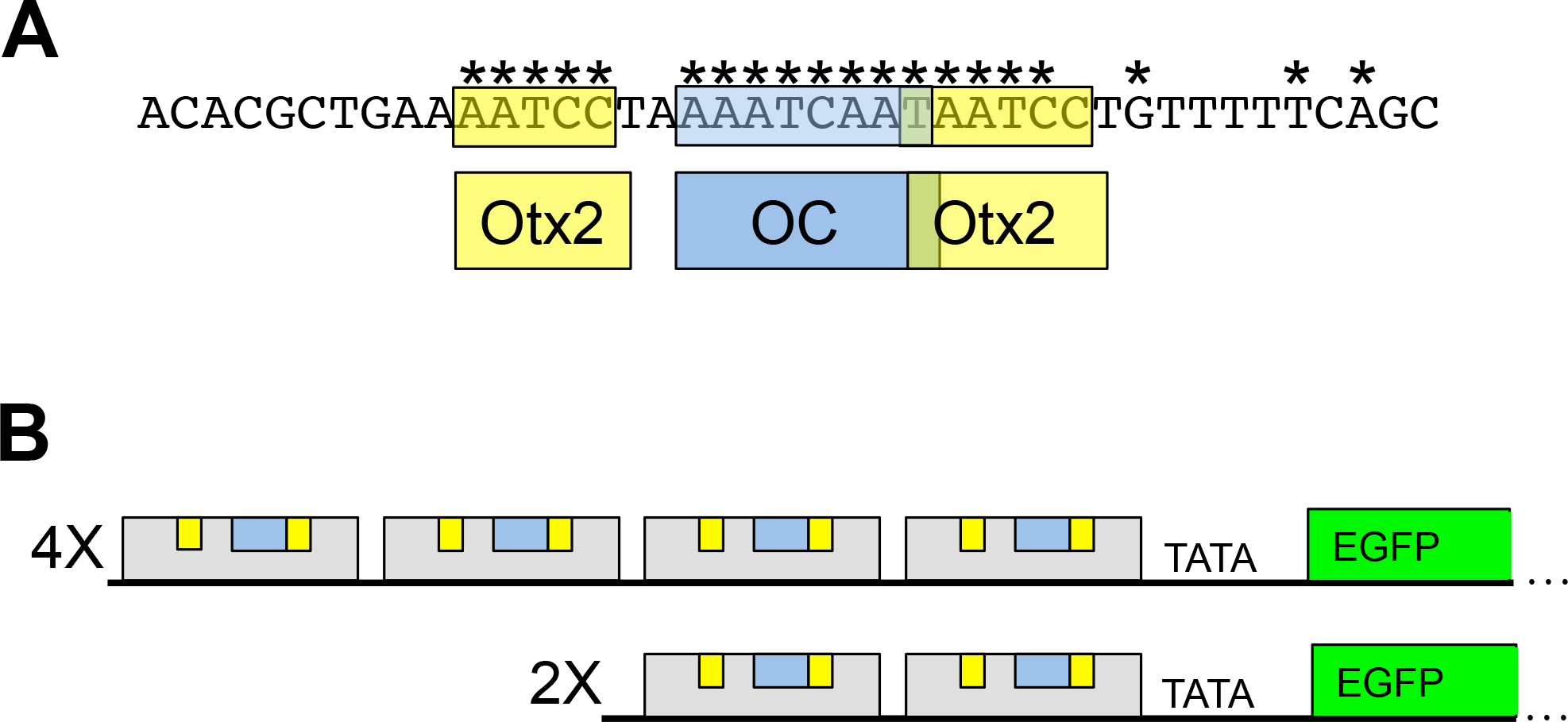
Schematic of the ThrbCRM1 Reporter and Sequence Elements. (A) Sequence of one copy of the ThrbCRM1 element and the corresponding binding sites of Otx2 and Onecut1 (OC1). (*) indicates conservation as defined in Emerson et al. 2013. (B) Schematic representation of the ThrbCRM1 construct with 4X or 2X copies of the 40 base pair ThrbCRM1 element shown as the grey box. Predicted Otx2 binding sites are indicated in yellow, and the predicted OC1 binding site is indicated in blue.

**Figure 2.**
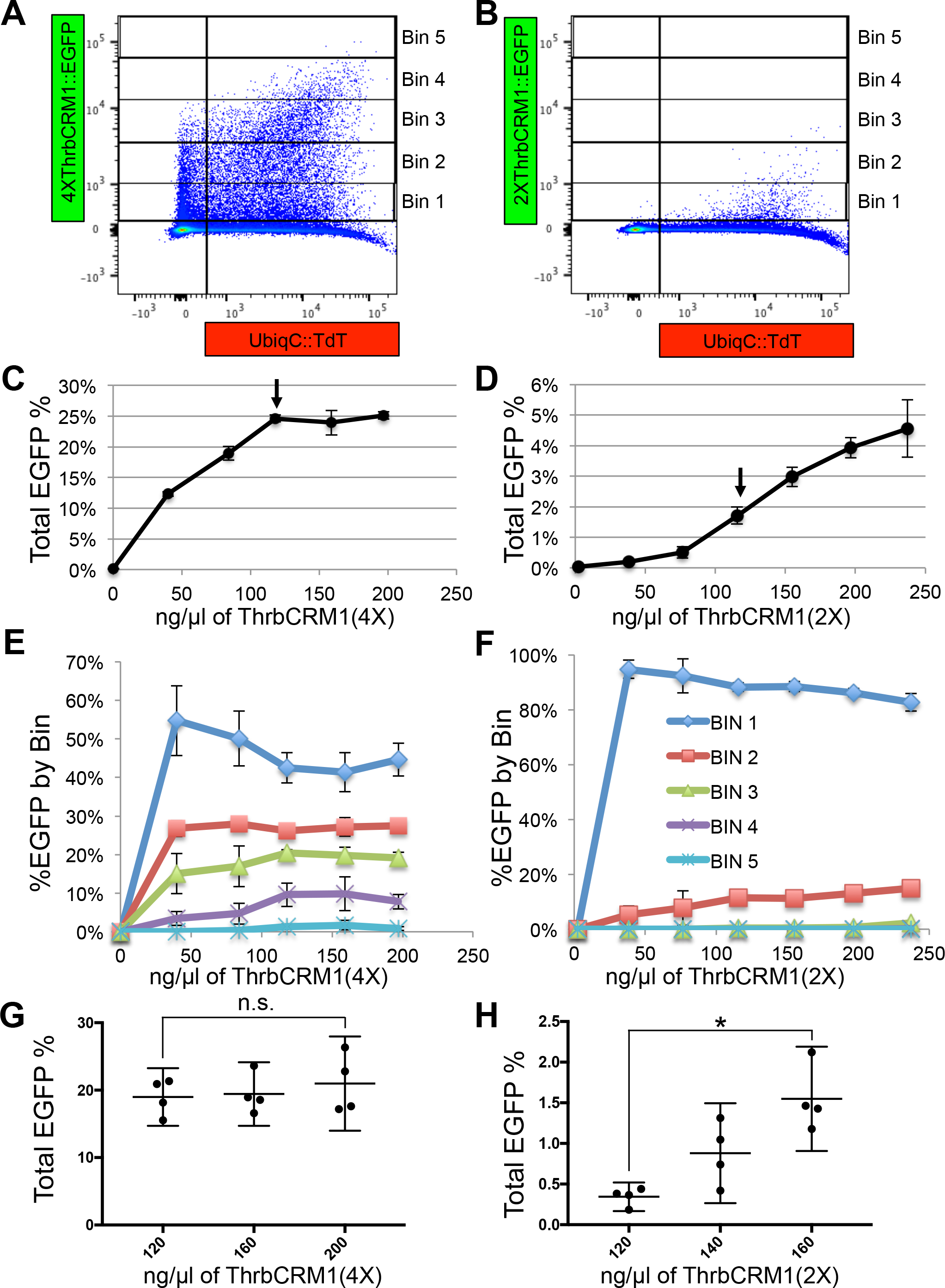
Effects of ThrbCRM1::GFP Plasmid Concentration and ThrbCRM1 Element Copy Number on Reporter Activity. (A,B) Representative flow cytometry plots of dissociated cells from chick retinas receiving 160 ng/μl of either the ThrbCRM1(4X) (A) or ThrbCRM1(2X) (B) EGFP reporter plasmid. Bins representing levels of EGFP fluorescence intensity are shown as vertical boxes and denoted on the right side of the plot. (C,D) Graphs of the percentage of total percentage of EGFP-positive cells along the y-axis relative to the concentration of the 4X (C) or 2X (D) ThrbCRM1::EGFP reporter plasmid on the x-axis. Arrow denotes 120ng/μl of reporter plasmid (E,F) A graph of the percentage of EGFP-positive cells in each bin out of the total number of EGFP-positive cells along the y-axis and the concentration of the reporter plasmid shown along the x-axis. Bin 1 through bin 5 represent increasing amount of EGFP fluorescence (bin 1 = least amount of EGFP fluorescence; bin 1 = dark blue; bin 2 = red; bin 3 = light green; bin 4 = purple; bin 5 = aqua) Bin key in panel F also applies to panel E (G,H) Graphs of the percentage of total EGFP-positive cells along the y-axis relative to the concentration of the 4X (G) or 2X (H) ThrbCRM1::EGFP reporter plasmid. Samples from G and H were generated in a single experiment, but plotted in separate graphs. Error bars represent 95% confidence intervals. Asterisk identifies a statistically significant p-value<0.5 using the Mann-Whitney t-test. n.s. denotes no significance.

**Figure 3.**
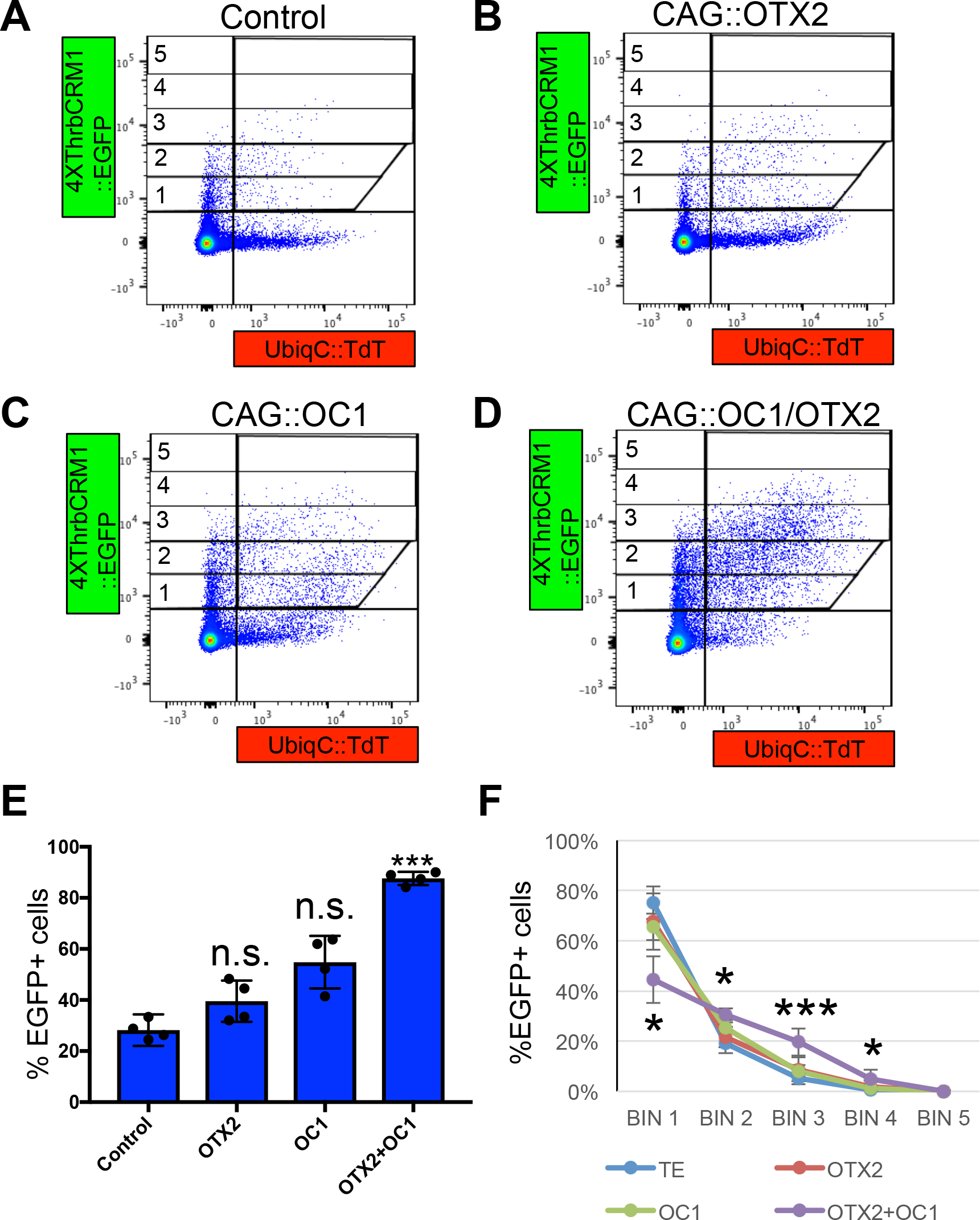
Effects of misexpression of Otx2 and/or OC1 on ThrbCRM1::EGFP Activity. (A-D) Representative flow cytometry plots of dissociated cells from chicken retinas receiving Ubiq::TdT reporter plasmid, 4XThrbCRM1::EGFP reporter plasmid and either TE (no DNA) (A), CAG::Otx2 (B), CAG::OC1 (C) or CAG::OC1 and CAG::Otx2 (D). (E) A plot of the percentage of ThrbCRM1::EGFP-positive cells in response to electroporation of the CAG plasmid shown along the x-axis. Average values are based on 4 retinas. (F) A graph of the percentage of ThrbCRM1::EGFP-positive cells in each bin (bins 1-5 as shown in A with bin 1 = least amount of EGFP fluorescence) for each of the 4 conditions. Plotted values represent the averages of 4 retinas and error bars represent 95% confidence intervals. A one-way Anova with a post-hoc Dunnetts statistical test was used to compare each of the misexpression groups to that of the TE group. * represents p<0.05, ** represents p<0.01 and ***represents p<0.001.

**Figure 4.**
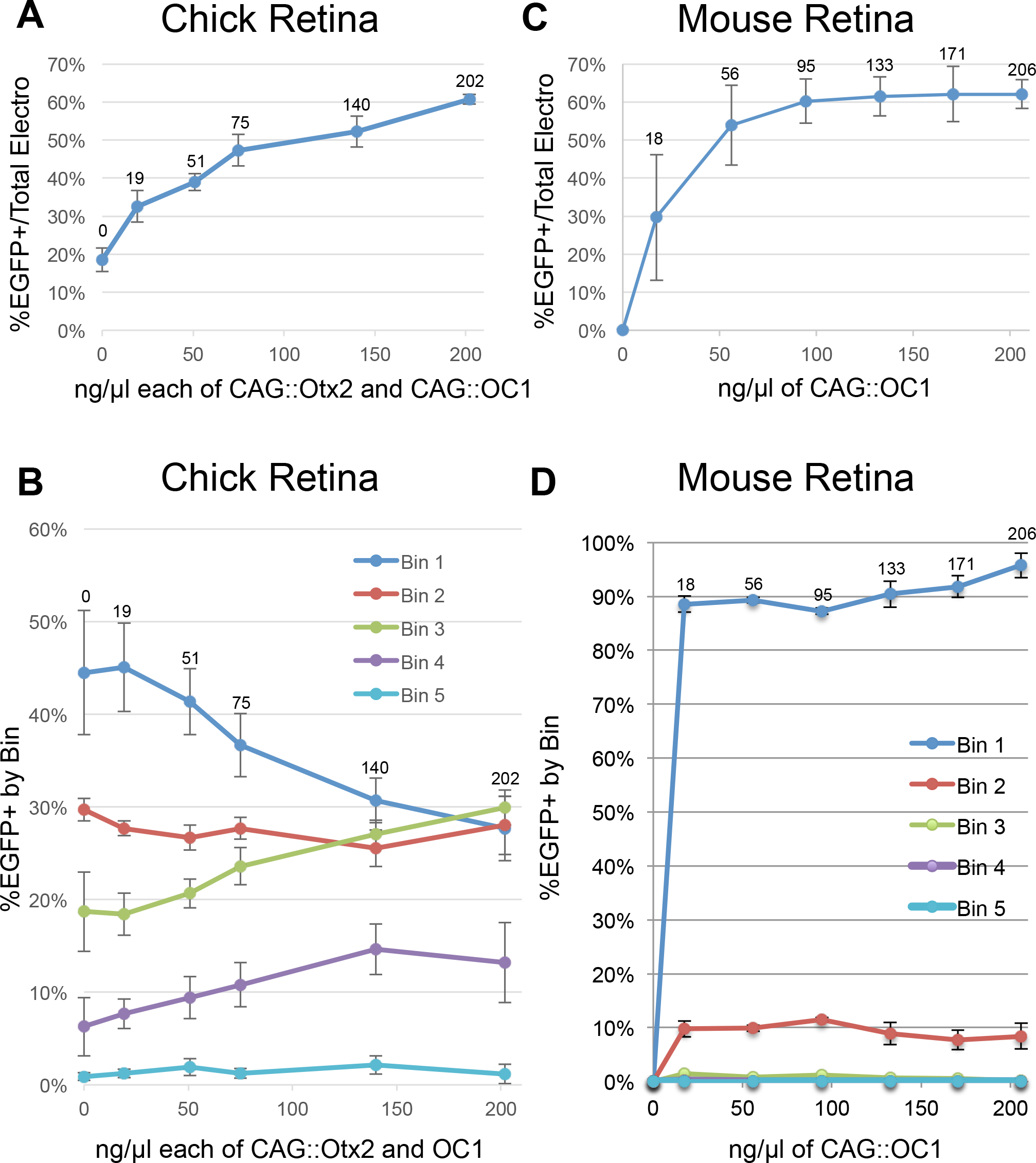
Effects of Transcription Factor Misexpression Plasmid Concentration on Reporter Activity in Mouse and Chick Retinas. (A) A graph of the percentage of electroporated cells in chicken E5 retina that are EGFP-positive after introduction of 160 ng/μl ThrbCRM1::EGFP reporter plasmid and varying concentrations of the CAG::Otx2 and CAG::OC1 misexpression plasmids. The x-axis displays concentrations in nanograms/microliter (ng/μl) of each of the misexpression plasmids. (B) A graph of the percentage of EGFP-positive cells in each bin (bins 1-5). Bin 1 through bin 5 represent increasing amount of EGFP fluorescence (bin 1 = least amount of EGFP fluorescence and bin 5 = the most amount of EGFP fluorescence). (C) A graph of the amount of cells positive for EGFP in a mouse P0 retina electroporated with 200 ng/μl of ThrbCRM1::EGFP reporter plasmid and varying concentrations of the CAG::OC1 misexpression plasmids. The x-axis displays concentrations in nanograms/microliter (ng/μl) of CAG::OC1 plasmid. (D) A graph of the amount of cells in each bin (bins 1-5) of the data plotted in C.

**Figure 5.**
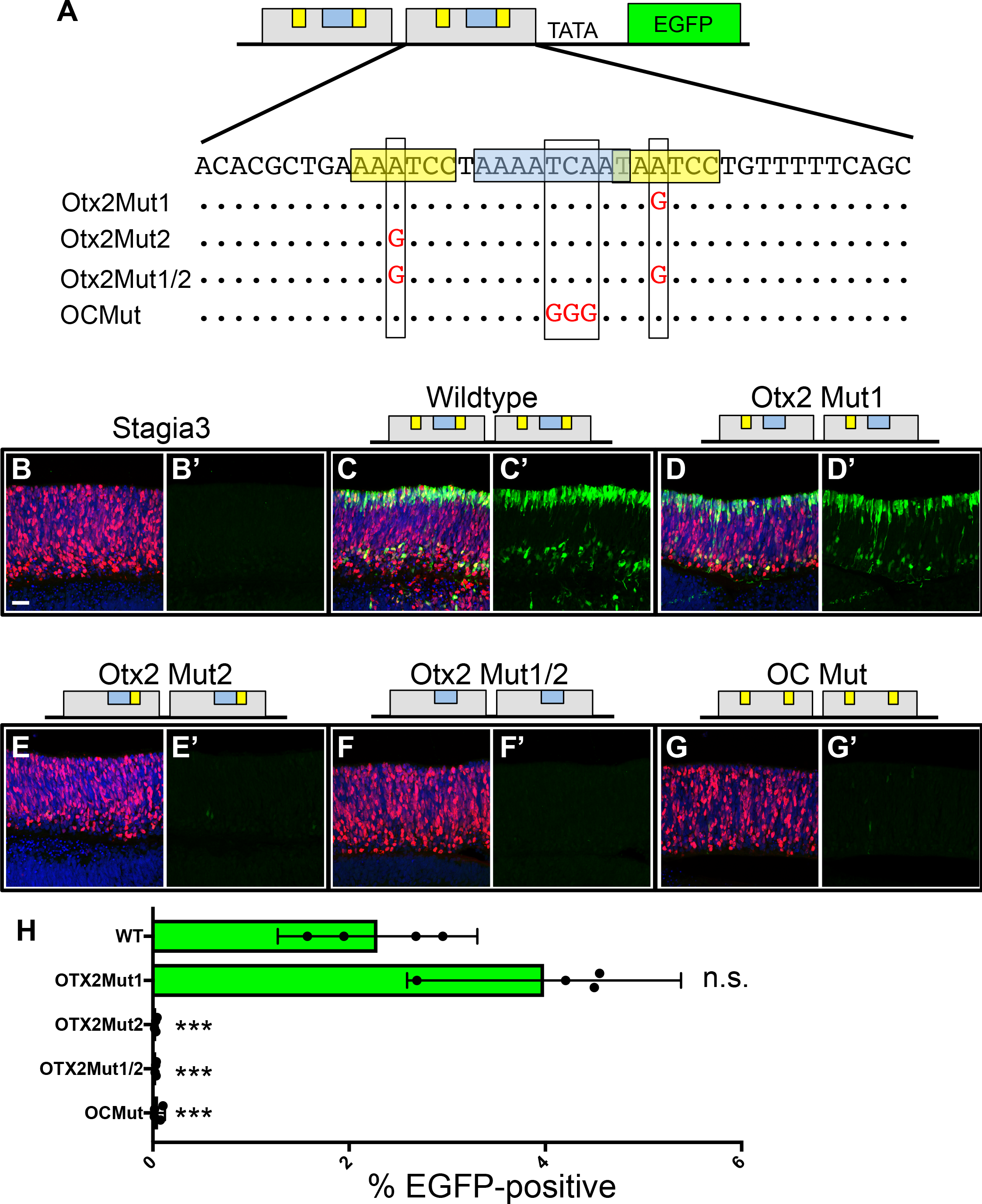
Mutational Analysis of the ThrbCRM1 Element. (**A**) Schematic and sequence representation of the ThrbCRM1 element. The putative OTX2 binding sites are highlighted in yellow and the OC1 binding site is highlighted in blue. The letters in red represent the mutation of the corresponding nucleotide. (**B**-**G, ‘B-G’**) Confocal z-stack images of chicken retinas electroporated with CAG::Nucβ-gal and the ThrbCRM1::EGFP plasmid shown above and immunofluorescent detection of EGFP (green), Nucβ-gal (red) and DAPI (blue). (**B-G**) Merged images (**B’-G’**) EGFP signal only (**H**) Results of a flow cytometry evaluation of EGFP fluorescence displayed as a graph of the percentage of cells positive for the ThrbCRM1-driven EGFP reporter (x-axis). *** denotes statistical significance p<0.001 as determined by a one-way Anova with a post-hoc Dunnetts test. The electroporated Stagia3 reporter is shown to the left of each bar. Scale bar in panel B represents 20μm and applies to all image panels.

### DNA electroporation mixes

DNA electroporation mixes were made with a volume of either 50 or 55ul, with 50ul used in the electroporation chamber for all experiments. A 10X phosphate buffered solution (PBS) was used to generate a final concentration of 1X PBS. To aid in accurate pipetting of viscous DNA solutions, a positive displacement pipettor (Eppendorf, Biomaster-4830) was used. For all experiments that involved a comparison between the effect of a particular plasmid, a mastermix was generated that included PBS and other plasmids found in all samples. For experiments in which the amount of EGFP reporter varied, 55ul DNA mixes were prepared by adding the determined amount of EGFP plasmid to a tube and preparing a mastermix of PBS and the other plasmids found in all samples before adding to the EGFP tubes. A Nanodrop blank sample was prepared by adding the appropriate volume of TE and mastermix and then prepared DNA mixes were measured at 260nm. The average of three spectrophotometer readings were used to empirically determine the amount of EGFP plasmid present in mixes, to avoid data skewing by pipetting error and was used in the plotting of data.

### DNA electroporations

Methodology for DNA electroporations was as described in (Emerson and Cepko, 2011) with the exception that a Nepagene Super Electroporator NEPA21 Type II was used to generate voltage pulses. Retinas were cultured for 2 days. For all electroporation experiments that used an EGFP reporter and a TdTomato co-electroporation control for flow cytometry analysis, a set of retinas were electroporated either with CAG::EGFP or UbiqC::TdT, and together with an unelectroporated retina, were used to generate compensation controls for the flow cytometer.

### Retina Dissociation

Retinal pigment epithelium and excess vitreal tissue was removed in HBSS media (GIBCO, 14170112) using forceps and retina was placed in microcentrifuge tube with 200μl HBSS. A papain activating solution of 200 μl/retina containing 11.6mM L-cysteine, 1.11mM EDTA and 5μl papain (Worthington Biochemical, L5003126) was added and incubated at 37°C for 15-25 minutes for chicken retinas or 35-45 minutes for mouse retinas. During this period, each tube was individually flicked to help break down the tissue into smaller clumps. 600 μl of 10%FBS (ThermoFisher, A3160602)/DMEM (Life Technologies, 11995-073) was added to stop the reaction. 10 μl/retina of DNase (Sigma-Aldrich 4536282001) was added, incubated in a 37°C water bath for five minutes, washed in DMEM and then fixed in 4% paraformaldehyde/1X PBS for 15 minutes. Cells were washed three times in 1 ml of 1X PBS upon being filtered through a 40μm strainer (Biologix, 15-1040). All centrifuge spins were at 1,700rcf for 5 minutes and supernatant removed with a P1000 pipettor.

### Flow cytometry

The dissociated single-cell suspensions were analyzed via flow cytometry using a BD Biosciences LSR II or FACS Aria machine. Approximately 300,000 cells were analyzed for each sample.

### Data quantitation and representation

Flow cytometry data was analyzed and plotted using Flowjo software. All experiments where percentages of EGFP-positive cells were calculated represent the averages calculated from 3 or 4 independently electroporated retinas. All average values refer to means and error bars in figures represent 95% confidence intervals. For plotting of Bin percentages in Figures 2 and 4, y-axis percentages were automatically set to “0” for samples in which no GFP reporter plasmid was added to prevent plot skewing by small numbers of cells. In cases where results were tested for statistical significance, a Mann-Whitney t-test was applied using JASP software(JASP Team, 2018). In cases in which a t-test was not appropriate because more than two groups were being compared, a one-way ANOVA with a post hoc Dunnetts test was applied using R 3.3.0. and the multcomp package (Hothorn et al., 2008; Team, 2018). All experiments were independently replicated and statistically analyzed to verify statistical significance of presented results.

### Immunofluorescence and Confocal Microscopy

Retinas analyzed for confocal microscopy were removed from filters after two days and fixed in 4% paraformaldehyde for 30 minutes at room temperature with gentle shaking in 24 well plates. After three washes with 1X PBS, retinas were sunk in 30%sucrose/0.5XPBS at 4 °C. Retinas were frozen in OCT (Sakura Tissue-Tek, 4583) and sectioned to 20μm thickness on a Leica CM1950 cryostat and placed on glass slides (FisherScientific, 12-550-15). Slides were processed for immunofluorescence as previously described (Emerson and Cepko, 2011). Primary antibodies used were chicken anti-GFP (abcam, 13970, 1:2,000) and mouse anti-β-galactosidase (DSHB, 40-1a-s, 1:20). Secondary antibodies were Goat anti-chicken Alexa488 (Jackson Immunoresearch, 103-545-155, 1:800) and Goat anti-mouse Cy3 (Jackson Immunoresearch, 115-165-146, 1:500). 4’,6-Diamidino-2-Phenylindole (DAPI) was applied in the third wash of PBT (1XPBS +0.1%Tween-20) at a final concentration of 1ug/μl. Slides were mounted in Fluoromount-G (Southern Biotech, 0100-01) with 34 X 60 cover slips (VWR, 48393 106) and sealed with nail polish (Sally Hansen 30003298000). Confocal images were acquired with a Zeiss 710 confocal using Zen Software (Zeiss, Version 2.1 Black 2015) and processed using FIJI 2/Image J software (Version 2.0.0-rc-67/1.52c).

## Results

### Experimental Paradigm Overview

To assess reporter activity in the retina, developing retinas from either chicken or mouse (Fig. 4C,D only) were isolated and electroporated with a DNA plasmid solution. Retinas were cultured ex vivo for 2 days, dissociated into single cells, and analyzed by flow cytometry. In all experiments, the activity of a cell-type specific EGFP reporter construct was assessed relative to a broadly active red fluorescent protein reporter construct. For both model animal systems, the ThrbCRM1 element was the main reporter plasmid used. ThrbCRM1 is a 40 base pair element containing a OC1 binding site and two potential Otx2 binding sites (Fig. 1A)(Emerson et al., 2013). Two different forms of the vector were used - “4X” and “2X” versions that contained four and two copies of the ThrbCRM1 element, respectively (Fig. 1B). The 4X version has a much greater overall activity level, likely due to the fact that there are twice as many binding sites for activating transcription factors and also that the four copies positions these binding sites further away from the basal promoter site, which may promote DNA looping regulatory events.

### Basal Vector Activity

The Stagia3 plasmid was used for all experiments and has been reported to have low basal activity in qualitative assessments (Billings et al., 2010; Blixt and Hallböök, 2016; Emerson and Cepko, 2011; Wang et al., 2014). A flow cytometry assay was used to quantitatively assess the basal activity of the Stagia3 plasmid. In addition, to determine if the presence of other plasmids with strong cis-regulatory elements (the CAG promoter element in this instance) could trans-activate the Stagia3 reporter, an increasing concentration of a CAG::Nuclear β-galactosidase (CAG::Nucβ-gal) plasmid was included. A third plasmid, CAG::mCherry was included at a constant concentration to standardize for the number of cells targeted by electroporation. Representative flow cytometry plots of retinas without any included CAG::Nucβ-gal plasmid (Fig. S1A) or 200ng/μl CAG::Nucβ-gal (Fig. S1B) show a large number of mCherry single-positive cells and very few cells that express EGFP. A plot of this data reveals that the basal vector used at the standard concentration of our ex vivo chicken experiments (160 ng/μl) had extremely low levels of EGFP expression in all samples. There was no statistically significant ectopic activation of EGFP even at the highest concentration of CAG-containing plasmids (Fig. S1C). This suggests that the Stagia3 reporter vector used in these experiments 1) possesses low basal activity 2) this basal activity is not altered by the presence of high concentrations of additional plasmids that possess strong regulatory elements.

### GFP reporter plasmid dose response curves

The Stagia3 reporter plasmid has been used in a number of studies to test the activity of potential cis-regulatory elements. In previous studies, a specific concentration (100ng/μl −200ng/μl, depending on the study) has been repeatedly used in chicken and mouse retinas, though it has not been empirically determined whether this is the ideal concentration for assessment (Billings et al., 2010; Emerson et al., 2013; Emerson and Cepko, 2011; Mo et al., 2016; Wang et al., 2014). When considering the electroporation of reporter plasmids, it is likely that three variables affect the amount of EGFP expression: the number of cells targeted (the number of cells that take up plasmids), the number of plasmids incorporated into each individual cell, and whether the transcription factors that regulate the cis-regulatory element are present in limiting amounts. We first sought to quantitatively measure the first two variables by determining the number of cells in the electroporated population that have any detectable EGFP expression and also to measure the relative fluorescence levels of the EGFP-positive population. Chicken E5 retinas were electroporated with concentrations between 0 and 200ng/μl of either the 4X or 2X ThrbCRM1::EGFP reporters and a fixed concentration (100ng/μl) of a broadly expressed TdTomato construct (UbiqC::TdTomato, hereafter referred to as UbiqC::TdT) to use as an independent measure of electroporation efficiency. Depicted in Fig. 2A and B are examples of the distribution of cells when 160ng/μl of either the 4X or 2X versions of the ThrbCRM1::EGFP reporter is used. The percentage of cells out of the entire electroporated population that were EGFP-positive was plotted against the concentration of reporter plasmid (Fig. 2C,D). It was observed that for the 4X plasmid, the number of EGFP-positive cells increased logarithmically and reached an asymptote of 25% of the entire electroporated population at a concentration of 120ng/μl of the reporter plasmid. In contrast to the ThrbCRM1(4X) element, the ThrbCRM1(2X) element was approximately 10-fold less active than the ThrbCRM1(4X) element and a plot of the concentration curve revealed a sigmoidal instead of hyperbolic shape. In addition, the percentage of EGFP-positive cells continued to rise at concentrations of reporter plasmid beyond 120ng/μl, and a clear plateau point was not observable for the concentrations tested. The observation that the 2X version was not saturating at the same concentrations as the 4X version suggests that the plateau effect observed for the 4X version is not simply a physical limit of the system, such as how much DNA can be electroporated. In addition, the plateau effect observed with ThrbCRM1(4X) suggests that there is a limit of 25% of cells at this time in development that can activate the ThrbCRM1 element. This is likely due to the number of cells co-expressing Otx2 and OC1.

In addition to calculating the total number of cells, the fluorescence intensity of the cells in that population was assessed by determining the distribution of the EGFP-positive cells across 5 Bins, with Bin1 being the weakest EGFP-expressing cells and Bin 5, the strongest expressing cells (Bin locations displayed in Fig. 2A,B). The distribution of cells relative to these Bins also stabilized at approximately 120ng/μl for the 4X version. At lower concentrations of reporter plasmids, most cells were located in the weakest EGFP-expressing Bin, which was Bin1. As the concentration of plasmids rose, there was an increase in the percentage of EGFP-positive cells in Bins 2, 3, and 4 and a subsequent decrease in the percentage of cells in Bin 1. The stabilization of these percentages suggest that there is a concentration threshold for the ThrbCRM1(4X)::EGFP reporter, such that the presence of more DNA plasmids in the electroporation mix does not lead to more cells activating the enhancer or for any of the cells to express more EGFP. Retinal cells electroporated with the 2X version expressed strikingly less EGFP compared to the 4X version. Almost no cells were found in Bins 3, 4, and 5 in the 2X version, whereas 18% and 8% of EGFP-positive cells were found with the 4X version.

The concentration curves for ThrbCRM1(X4) and ThrbCRM1(X2) shown in Fig. 2 (C and D) were generated in separate experiments, which can lead to experiment-specific differences based in embryo timing or flow cytometer settings. To confirm that the ThrbCRM1(X4) and ThrbCRM1(X2) constructs saturated at different points, the two constructs were tested in the same experiment across a range of concentrations for which the (X4) was saturated and the (X2) was not (120, 160, 200 ng/μl) (plotted separately in Fig. 2 G,H to more easily allow for comparison of concentration differences irrespective of scale). Indeed, the concentration-dependent differences were observed and a comparison of the EGFP reporter activity of the 120 ng/μl and the 200 ng/μl concentrations of each plasmid was statistically different for the 2X version but not the 4X version. This confirms that the presence of 4 sets of binding sequences compared to 2 sets of binding sequences not only results in a larger EGFP-positive population with individual cells expressing more EGFP, but that the saturation points for these metrics are shifted to lower concentrations.

### Effects of misexpression of Otx2 and OC1 on the ThrbCRM1 population

A previous study has identified the ThrbCRM1 element as containing predicted Onecut and Otx2 binding sites and a chromatin immunoprecipitation experiment confirmed their occupancy of this element in the developing chicken retina (Emerson et al., 2013). The current model is that these two transcription factors are co-expressed in a subset of retinal progenitor cells and both transcription factors are required for co-activation of ThrbCRM1 element-driven reporter expression (Emerson et al., 2013). Each of these transcription factors is also expressed without the other one in certain populations of retinal cells, including retinal progenitor cells (Buenaventura et al. 2018). To determine whether the population of retinal cells that activate the ThrbCRM1 element could be expanded, an experiment was performed in which retinas were co-electroporated with the ThrbCRM1(4X)::EGFP reporter, a UbiqC::TdT co-electroporation control, and plasmids that drive the broad expression of Otx2 and/or OC1 transcription factors (using the CAG promoter). When either the Otx2 or OC1 misexpression plasmids were introduced, an increase in the proportion of the electroporated population that activated the ThrbCRM1(4X)::EGFP reporter was observed, though these increases were not statistically significant (Fig. 3A-C). Inclusion of both the Otx2 and OC1 misexpression plasmids led to a statistically significant increase in the EGFP population (Fig. 3D). The percentage of the electroporated population that activates the ThrbCRM1 element under these conditions is plotted (Fig. 3E). One interpretation of these results, based on the current model of ThrbCRM1 activation, is that misexpression of only one of the transcription factors, for instance just Otx2, expands the population of cells that activate ThrbCRM1 to those cells that normally only express OC1. The same would be true when OC1 is misexpressed, as normally Otx2-only cells would now activate the ThrbCRM1 element. Misexpression of both transcription factors leads to additional activation of the ThrbCRM1 element in cells that do not normally express either transcription factor. While the percentage of cells that activate ThrbCRM1 in response to the inclusion of both transcription factors increases dramatically, it does not lead to EGFP expression in all of the electroporated population. This could reflect a technical limitation of co-electroporation efficiency of all of these plasmids or it could reveal a biological limitation. Perhaps there are repressive transcription factors expressed in a subpopulation of cells that interact with ThrbCRM1 to keep it off even in the presence of Otx2 and OC1. Alternatively, there may be differentially expressed cofactors that are necessary to cooperate with Otx2 and OC1 to activate the ThrbCRM1 element.

The distribution of EGFP-positive cells in these transcription factor misexpression experiments was calculated using the same Bin system as described above in Fig. 2 (Fig. 3F). Interestingly, in retinas that had just Otx2 or OC1 misexpressed, the distribution of EGFP-positive cells across the five Bins was not statistically different from that of retinas with just the ThrbCRM1 reporter introduced. In contrast, in retinas with both transcription factors misexpressed, there was a statistically significant redistribution of EGFP-positive cells to Bins containing higher EGFP fluorescence. One interpretation of this result is that in retinas in which only one transcription factor was misexpressed, the new population of cells that activates ThrbCRM1 may be limited by the amount of the endogenous transcription factor that is present. Thus, while these cells may ectopically express the reporter, their fluorescence intensity would be similar to the cells that normally activate the reporter. However, in the case where both Otx2 and OC1 are introduced, these two proteins are now both present at higher concentrations than their endogenous levels and this leads to a higher amount of EGFP production per cell that is quantitatively captured through this Bin analysis.

To determine the concentration effect of misexpressing the Otx2 and OC1 factors on ThrbCRM1(X4) activity, plasmids encoding misexpression constructs for both transcription factors were introduced at similar relative proportions to each other. A dose-dependent increase in activity was observed with a distribution that suggested that even one-tenth the amounts of misexpressed Otx2 and OC1 that were used previously were sufficient to increase the output of reporter activity (Fig. 4A). While a clear plateau point in the total GFP-positive population was not observed by the highest amount of DNA tested (202 ng/μl), the bin distribution of EGFP fluorescent cells displayed a plateau beginning at approximately 75 ng/μl (Fig. 4A,B).

### ThrbCRM1 Activity in the Mouse postnatal retina

The ThrbCRM1 element is not active when electroporated into the mouse postnatal retina (Emerson et al., 2013). This is due to the fact that at this time, Onecut family members are only expressed in postmitotic cells, which are not targeted by electroporation in this paradigm. However, Otx2-positive cells can be targeted and co-electroporation of a CAG::OC1 misexpression plasmid with ThrbCRM1::EGFP leads to robust upregulation of EGFP reporter activity as well as upregulation of endogenous gene expression associated with cones and horizontal cells (Emerson et al., 2013). In support of the requirement of Otx2, simultaneous removal of Otx2 via a floxed allele leads to a concomitant decrease in cells with positive ThrbCRM1 activity (Emerson et al., 2013). These experiments did not determine the concentration requirements for OC1 and so an experiment to do so was designed. Postnatal day 0 (P0) mouse retinas were electroporated with a constant level of the ThrbCRM1(4X)::EGFP reporter and a UbiqC::TdT construct. A variable amount of CAG::OC1 was co-electroporated. A plot of this data revealed a CAG::OC1 concentration-dependent curve with a hyperbolic form that plateaued between 56ng/μl and 95ng/μl (Fig. 4C). Analysis of the EGFP intensity distribution of cells across Bins did not reveal major differences between the lowest effective concentration (18ng/μl) and the highest (206ng/μl) (Fig. 4D). This suggests that the previously used concentration of 100ng/μl was near the plateau point, but also reveals that much lower concentrations of the misexpression plasmid are biologically active and could be used to examine biological effects on cell fate.

### Mutational analysis of the ThrbCRM1 element

A previous study identified OC1 and Otx2 binding sites in the ThrbCRM1 element through chromatin immunoprecipiation and functionally tested the requirement of the OC1 binding site by mutating 5 of the core base pairs that compose this site (Emerson et al., 2013). However, only a qualitative readout was used to assess whether the OC1 binding site mutation affected the activity of the element and whether either of the two Otx2 binding sites was required was not determined. To examine the DNA sequence requirements more closely, specific point mutations were introduced to disrupt the putative Otx2 binding sites either alone (Otx2Mut1 or Otx2Mut2) or together (Otx2Mut1/2) (Fig. 5A). For both potential binding sites, the second Adenine of the motif was mutated as a recent high-throughput Selex study identified an Adenine in this position in all recovered Otx2 bound sequences (Jolma et al., 2013). A separate 3 base pair mutation was made in the predicted OC1 binding site (OCMut) (Fig. 5A). These mutations were made in the context of the ThrbCRM1(2X)::EGFP reporter where it was possible to efficiently introduce identical mutations into both copies of the ThrbCRM1 element.

We first qualitatively tested these constructs in the context of the intact retina (Fig. 5B-G). ThrbCRM1(2X)::EGFP constructs were co-electroporated with a CAG::Nucβ-gal construct to identify electroporated areas of the retina. Retinas were examined for EGFP and Nucβ-gal reporter activity using confocal microscopy. As expected, a Stagia3 plasmid without additional cis-regulatory elements was unable to drive EGFP expression (Fig. 5B, B’). The wild-type construct showed a previously characterized pattern indicative of apically located photoreceptors, possible ThrbCRM1-active retinal progenitor cells and basally located horizontal cells while the Nucβ-gal was found throughout the retinal thickness (Fig. 5C, C’). Surprisingly, the number of EGFP-positive cells in the Otx2Mut1 electroporated retina was similar to that observed with the wildtype construct (Fig. 5D, D’). This sequence is highly conserved in vertebrates and matched the Otx2 binding site the most closely out of the two potential Otx2 binding sites. In contrast, very few EGFP-expressing cells were observed in retinas electroporated with the Otx2Mut2, Otx2Mut1/2 and OCMut constructs compared to the retinas with the WT and Otx2Mut1 constructs (Fig. 5E-G’). These results suggest that the sequence mutated in the Otx2Mut2 region, and not the Otx2Mut1 region, is the site that Otx2 binds in the ThrbCRM1 element.

To more quantitatively examine the effect of these mutations, each ThrbCRM1(2X)::EGFP construct was co-electroporated with UbiqC::TdT and analyzed by flow cytometry. The percentage of cells that activated the EGFP reporter was calculated and plotted (Fig. 5H). The wildtype version of ThrbCRM1(2X)::EGFP activated the reporter in just over 2% of the electroporated cells, in agreement with the low number of EGFP-expressing cells predicted from the concentration-dependent curves. In accord with the confocal microscopy results, the Otx2Mut1 reporter construct did not have a decrease in EGFP activity, and in fact had a slight increase. In contrast, mutation of the other potential Otx2 binding site (Otx2Mut2) led to a total abrogation of reporter activity and a plasmid carrying both mutations (Otx2Mut1/2) similarly had no EGFP expression. A plasmid carrying a mutation predicted to disrupt the OC1 binding site had a significant reduction in EGFP activity. Taken together, these results identify the critical Otx2 and OC1 binding sites for ThrbCRM1 element activity necessary for ThrbCRM1-driven reporter activity using both qualitative and quantitative assays.

### Misexpression of OC1 or Otx2 can partially activate ThrbCRM1 mutant elements

To further test the requirements of the binding sites identified in the mutagenesis experiments, we determined whether misexpression of either OC1 or Otx2 could activate mutated ThrbCRM1 reporters that lacked OC1 or Otx2 binding sites. Mutated ThrbCRM1 reporters were electroporated into chicken E5 retinas in combination with Otx2 or OC1 misexpression plasmids and analyzed by flow cytometry. We first tested the ThrbCRM1[Otx21/2Mut]::EGFP plasmid, which lacks any consensus Otx2 binding sites (Fig. 6A). As expected, misexpression of Otx2 did not significantly activate EGFP expression from the ThrbCRM1[mutOtx1/2]::EGFP construct when compared to a control. Surprisingly, electroporation of CAG::OC1 with the ThrbCRM1[MutOtx1/2]::EGFP construct induced EGFP expression. This suggests that excess OC1 is able to activate the ThrbCRM1 element even under conditions in which consensus Otx2 binding sites are absent. We next tested whether mutation of the OC binding site would affect the ability of the OC1 and Otx2 misexpression plasmids to activate the ThrbCRM1::EGFP plasmid. Similarly, Otx2 misexpression was able to significantly increase the amount of EGFP-positive cells from the OC mutated ThrbCRM1 plasmid, while the OC1 plasmid was unable to do so (Fig. 5B). This suggests that excess Otx2 can activate the ThrbCRM1 plasmid even when OC1 consensus binding sites are lacking in the ThrbCRM1 element. Taken together, these results 1) provide further confirmation that the sites targeted for mutagenesis are in fact the relevant binding sites for their cognate transcription factors and 2) the mutated reporter plasmids can be activated under conditions in which the transcription factor with an intact binding site is misexpressed.

**Figure 6.**
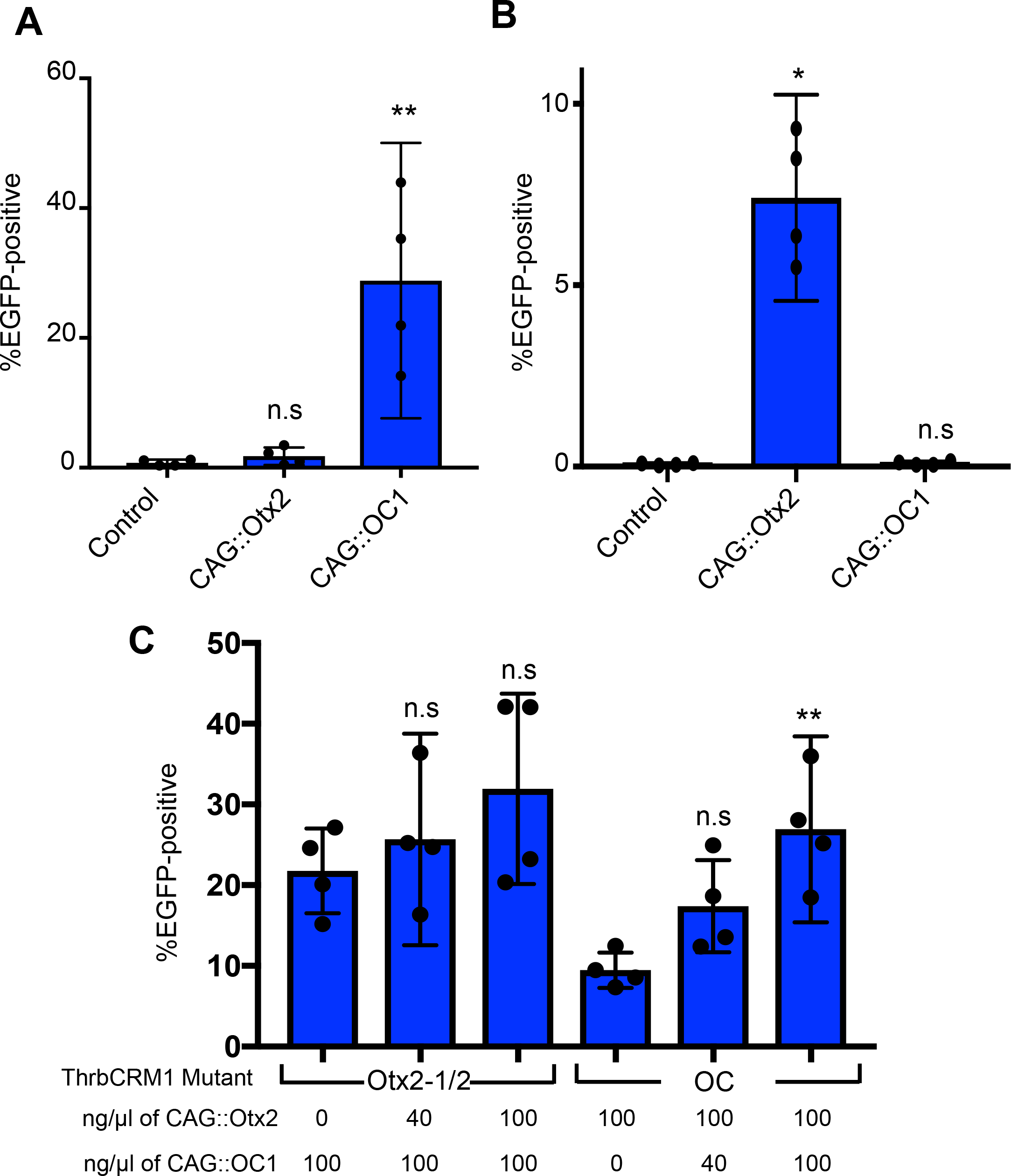
Activity of mutant ThrbCRM1::EGFP reporter in response to misexpression of Otx2 and/or OC1. (A-B) Graphs of the percentage of electroporated cells positive for EGFP after introduction into E5 chick retinas of 160 ng/μl of ThrbCRM::EGFP reporter plasmid with either no other plasmid, 100 ng/μl of Otx2 misexpression plasmid, or 100 ng/μl of OC1 misexpression plasmid. (C) A graph of the percentage of EGFP-positive cells (y-axis) in the electroporated population after electroporation of E5 chick retinas with the ThrbCRM1 mutant reporter plasmid and the concentration of the Otx2 and OC1 misexpression plasmids (ng/μl) shown along the x-axis for each condition. Error bars represent 95% confidence intervals. Statistical significance is denoted by *p<0.05 or **p<0.01 as determined by a one-way Anova with a post-hoc Dunnetts test.

These experiments suggest that the necessity for one of the transcription factor binding sites can be overcome by misexpression of the other transcription factor that has an intact binding site. Two major possibilities exist to explain this phenomenon. One is that under normal conditions, co-expression of both transcription factors is needed to lead to stable occupancy of ThrbCRM1 and detectable reporter expression. High misexpression of one of the transcription factors could lead to stable occupancy and reporter activation by this transcription factor, independent of the other transcription factor. However, this explanation is not supported by previous data in which misexpression of either Otx2 or OC1 in tissues that lacked the other transcription factor were unable to induce the ThrbCRM1 reporter (Emerson et al., 2013). A second possibility is that increased expression of one transcription factor may allow it to bind to its site, while recruiting the other transcription factor through a largely DNA-binding independent process. The lack of activity of ThrbCRM1 in tissues that only expressed one of the factors, no matter how highly expressed the other one, would be congruent with this hypothesis. To discriminate between these hypotheses, we repeated the previous misexpression experiments shown in Fig. 6A and 6B, but also included increasing concentrations of the misexpression plasmid encoding the other, presumptive non-DNA binding, transcription factor. In both cases, co-electroporation of the plasmid encoding the transcription factor that lacks a consensus binding site on the mutated ThrbCRM1 element led to a concentration-dependent increase in EGFP expression from the mutated ThrCRM1 reporter. In the case of the Otx2 mutant reporter, this was not a significant increase, while the OC mutant reporter was significant between the lowest and highest levels of CAG::OC1 tested. This suggests, that the OC1 transcription factor is able to participate in activation of the ThrbCRM1 element even under conditions where a consensus binding site is lacking.

## Discussion

The analysis of GRNs has yielded insights into fundamental developmental processes (Buecker and Wysocka, 2012; Peter and Davidson, 2016). However, cis-regulatory elements are a critical component of GRNs that have proven difficult to analyze at the quantitative level. Most studies, done in vertebrates, are limited to the identification of these elements and assessing the effects of mutations through qualitative assays, though quantitative analysis through fluorescence measurements of whole tissue has been reported (Montana et al., 2013). Investigating the nature of interactions between DNA elements and transcription factors is often limited to in vitro assays where differences in binding partners and cellular context are lacking. Thus, the generation and use of quantitative assays in the context of developing tissue, as shown here, is of critical importance.

This study shows that the concentration of reporter plasmid is an important variable in experimental design, though this aspect has been ignored by the vast majority of previously published electroporation studies. In cases of sequence element mutagenesis, the ideal concentration of reporter plasmid is below the saturation point for reporter output. This concentration allows for the detection of either partial loss of activation or an increase in reporter activity due to loss of a repressor site. Use of reporter plasmid concentrations above the saturation point could obscure meaningful reporter output changes. This study also demonstrates that saturation points are likely to be unique for a given cis-regulatory element in a particular biological context. In addition to these important technical considerations, we also suggest that the quantitative flow cytometry assay used here reveals saturation kinetics that are a direct result of transcription factor occupancy of cis-regulatory elements. A confirmation of the direct binding kinetics of the transcription factors in these cells is not possible with current techniques, but the increased fluorescence levels at the per cell level induced by increased transcription factor expression supports this interpretation.

Similar to reporter plasmids, a systematic evaluation of misexpression plasmid concentration in electroporation experiments has not previously been examined. Given the demonstrated effects of transcription factor expression levels inducing specific cell fates (e.g. Gli levels in the spinal cord, or Otx2 levels in the retina)(Stamataki et al., 2005; Wang et al., 2014), the concentration of misexpression plasmids should be considered in experimental design. For example, misexpression of OC1 in the postnatal retina is able to induce the earliest known steps of cone genesis in developing retinal cells that do not normally generate cones (Emerson et al., 2013). However, these cells appear stalled in their cone differentiation program, perhaps as a result of high levels of sustained OC1 expression in these cells. The present work shows that significantly lower concentrations of OC1 are sufficient to induce the ThrbCRM1 element in the mouse retina. Thus, it will be of interest to assess whether these lower levels of misexpressed OC1 influence the progression of the cone differentiation program.

The small size of the ThrbCRM1 element and the known identity of two key regulators of its activity make this cis-regulatory element an ideal candidate to explore at the functional level. The limited sequence constrains the regulatory information that can be encoded and thus the complexity of its regulation. Otx2 and OC1 are the only known regulators and the current model supports a necessary and sufficient model for their regulation of the ThrbCRM1 element (Emerson et al., 2013). The mutagenesis experiments performed here further support this model as sequences corresponding to one binding site for each of the transcription factors were found to be required for ThrbCRM1 activity. We interpret the ineffectiveness of misexpression of the corresponding transcription factor in eliciting reporter activity as further evidence that these mutations prevent most, if not all, of Otx2 and OC1 binding. However, we cannot rule out a small amount of binding that is perhaps facilitated by the presence of the other transcription factor with an intact binding site. Thus, while the misexpression experiments in the context of mutated binding sites supports a model where OC1 is recruited through protein-protein interactions with Otx2 and modulates transcription irrespective of DNA-binding, this assay may reveal the weak binding affinity of OC1 for the mutated binding site. Regardless, the dose-dependence of this effect indicates a co-regulatory function for these two proteins. This could reflect a physical interaction between these proteins, which is likely given the proximity of the Otx2 and OC1 binding sites.

This paradigm allowed us to test the necessity of sequence elements in the ThrbCRM1 element. Interestingly, only one of the two potential Otx2 binding sites appeared to be necessary for the activity of the ThrbCRM1 element. Whether the spacing and orientation of this Otx2 site relative to the OC1 binding site is important will require further investigation. In addition to mutagenesis experiments for this purpose, it will be interesting to identify other cis-regulatory elements co-regulated by Otx2 and OC1 and examine their binding sites with regards to specific sequences, orientation, and spacing. The likely Otx2 binding site (AAATCC) differs from the canonical monomeric site identified in in vitro Selex studies (TAATCC)(Jolma et al., 2013). In this Selex study, sequences containing Adenine in the first position were recovered, suggesting that A can be tolerated by Otx2 binding. However, though an A represented the second most enriched base after T, this was only 3.3% of all sequences. Interestingly, the same Selex study identified presumed Otx2 dimer-bound sequences and the sequences that correlated to the individual units differed in sequence from those bound by the monomer. A similar divergence in monomer versus dimer-bound sequence specificities has also been suggested for the Otx2-related transcription factor Crx, implying that the binding specificities of the Otx2 class of transcription factors can differ depending on their interactions with DNA as a monomer or a dimer (Hughes et al., 2017; Kwasnieski et al., 2012). We speculate that the AAATCC sequence could represent the preferred sequence for Otx2 only when present in a complex with OC1. Such a shift in binding specificity has been referred to as “latent specificity” and observed previously for Drosphila Hox genes when in complex with the cofactor Extradenticle (Slattery et al., 2011). This could provide a potential explanation for the high degree of conservation of an A in the first position that is found in the homologous ThrbCRM1 elements across the phylum chordata.

In summary, this study quantitatively assesses the effects of multiple experimental parameters on the activity of the restricted RPC ThrbCRM1 element. This has provided insights not only into the specific sequence requirements and transcription factor/DNA binding dynamics for this particular element, but more generally into the use of electroporation to investigate cis-regulatory elements during development.

## Acknowledgements

The β-galactosidase antibody developed by Joshua Sanes was obtained from the Developmental Studies Hybridoma Bank developed under the auspices of the NICHD and maintained by the Department of Biology at The University of Iowa. We thank Andre Washington for excellent technical help in tissue processing. Jeffrey Walker and Jorge Morales provided valuable technical support with flow cytometry and confocal microscopy. Diego Buenaventura provided essential input on statistical methodology. We thank Julianna LeMieux for editing/proofreading the manuscript.

## Ethics approval and consent to participate

All animal procedures were approved by The City College of New York Institutional Animal Care and Use Committee under protocol 932.

---

**Supplemental Figure 1. Assessment of plasmid transactivation on Stagia3 reporter plasmids**. (**A-B**) Representative flow cytometry plots of dissociated cells from chicken retinas electroporated with an empty Stagia3 reporter plasmid, a CAG::mCherry co-electroporation control, and another plasmid containing either 0 ng/μl (A) or 200 ng/μl (B) of an additional plasmid containing the CAG element without a fluorescent readout. Stagia3 reporter activity is plotted along the y-axis and the CAG::mCherry co-electroporation control along the x-axis. (**C**) Quantification of Stagia3 reporter plasmid activity plotted along the y-axis as the percentage of EGFP-positive in the electroporated population in the presence of varying concentrations of CAG co-electroporated plasmid in nanograms/microliter along the x-axis. Error bars represent 95% confidence intervals. There was no statistically significant effect of any concentration of additional CAG plasmid concentration compared to the baseline as assessed by a one-way Anova with a post-hoc Dunnetts test.

