## Supplementary figures and images for "Quantitative analysis of the ThrbCRM1-centered gene regulatory network"

### Supplemental Figure 1

#### Supplemental figure 1

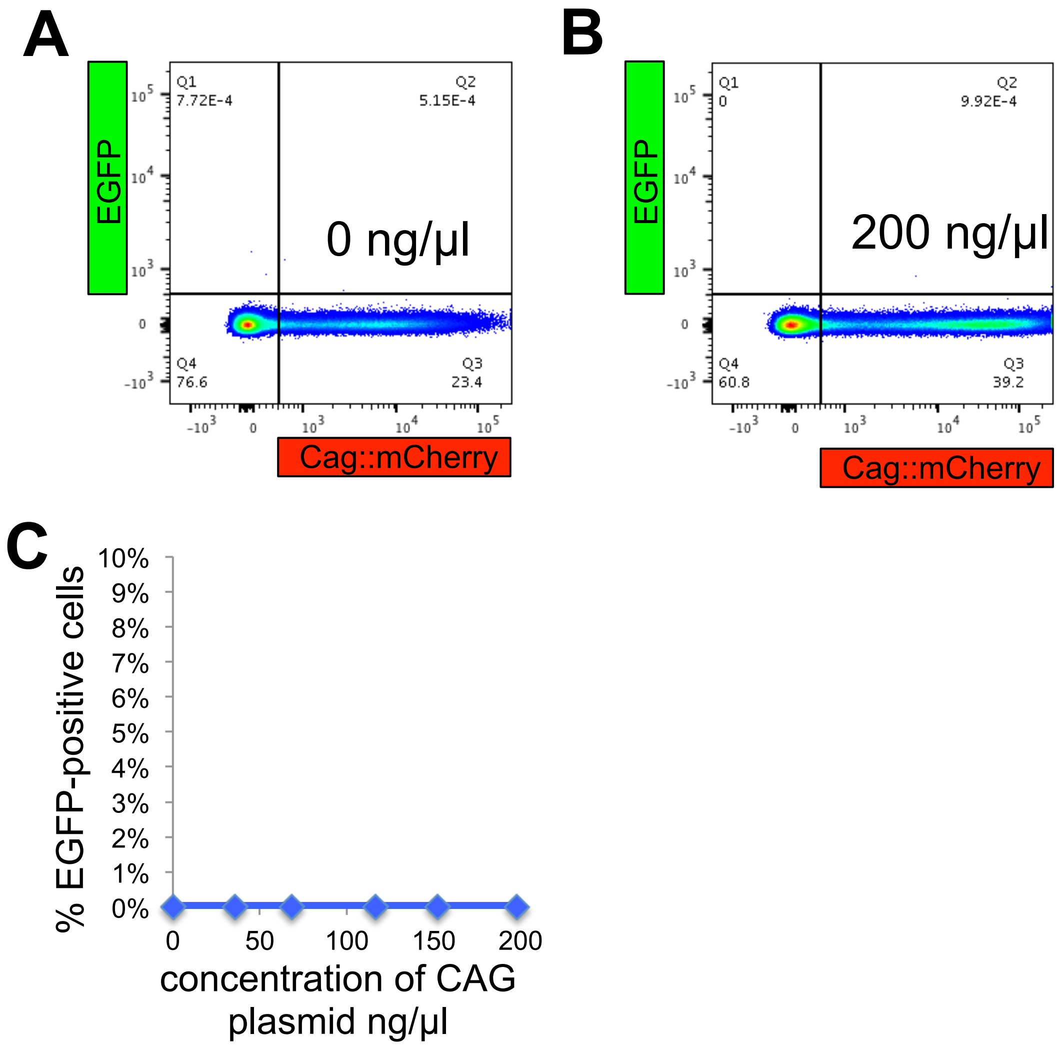
